## Supplemental Table 1 for "TRIM28 is a target for paramyxovirus V proteins"

**Supplementary Table 1. Primer pairs used for RT-qPCR analyses.**

| **Name** | **Fw/Rv** | **Sequence** |
| --- | --- | --- |
| **IAV M (WHO M30F2)** | Fw | ATGAGYCTTYTAACCGAGGTCGAAACG |
| **IAV M (WHO M264R3)** | Rv | TGGACAAANCGTCTACGCTGCAG |
| **PIV2 P** | Fw | TGCATCTTTTATAACTACTGATCTTGCTAA |
| **PIV2 P** | Rv | GTTCGAGCAAAATGGATTATGGT |
| **PIV5 M** | Fw | TCATGAGCCACTGGTGACAT |
| **PIV5 M** | Rv | TGGAATTCCCTCAGTTGTCC |
| **18S-rRNA** | Fw | GGCCCTGTAATTGGAATGACTC |
| **18S-rRNA** | Rv | CCAAGATCCAACTACGAGCTT |
| **ERV_9.1** | Fw | TCTTGGAGTCCTCACTCAAACTC |
| **ERV_9.1** | Rv | ACTGCTGCAACTACCCTTAAACA |
| **HERVK14C** | Fw | GTAATTGTGAGTACCCAAAATCTC |
| **HERVK14C** | Rv | ACCTTGTCCCAATCTTTTAC |
| **LTR13** | Fw | AAGCTGGCCCACAGTTATCC |
| **LTR13** | Rv | AGGTGCACAGAGTGGAACAG |
